# Sherlock - A flexible, low-resource tool for processing camera-trapping images

**DOI:** 10.1101/2023.03.08.531714

**Authors:** Matthew J. Penn, Verity Miles, Kelly L. Astley, Cally Ham, Rosie Woodroffe, Marcus Rowcliffe, Christl A. Donnelly

**Affiliations:** University of Oxford; The Zoological Society of London

## Abstract

1. The use of camera traps to study wildlife has increased markedly in the last two decades. Camera surveys typically produce large datasets which require processing to isolate images containing the species of interest. This is time consuming and costly, particularly if there are many empty images that can result from false triggers. Computer vision technology can assist with data processing, but existing artificial intelligence algorithms are limited by the requirement of a training dataset, which itself can be challenging to acquire. Furthermore, deep-learning methods often require powerful hardware and proficient coding skills.
2. We present Sherlock, a novel algorithm that can reduce the time required to process camera trap data by removing a large number of unwanted images. The code is adaptable, simple to use and requires minimal processing power.
3. We tested Sherlock on 240 596 camera trap images collected from 46 cameras placed in a range of habitats on farms in Cornwall, UK, and set the parameters to find European badgers (*Meles meles*). The algorithm correctly classified 91.9% of badger images and removed 49.3% of the unwanted ‘empty’ images. When testing model parameters, we found that faster processing times were achieved by reducing both the number of sampled pixels and ’bouncing’ attempts (the number of paths explored to identify a disturbance), with minimal implications for model sensitivity and specificity. When Sherlock was tested on two sites which contained no livestock in their images, its performance greatly improved and it removed 92.3% of the empty images.
4. Though further refinements may improve its performance, Sherlock is currently an accessible, simple and useful tool for processing camera trap data.

## 2 Introduction

Camera-trapping methods have the potential to provide profound insights into the behaviour and ecology of animal populations [1] and are becoming increasingly widespread across the world [2]. Development in technology has enabled the simultaneous study of a range of species across large spatial scales and with minimal disturbance. It is a substantially more cost-effective method of surveying mammals than the more traditional live trapping [3] and has led to the creation of extensive digital datasets of images [4] on a scale that was previously unachievable.

A common constraint of camera-trapping research is the vast quantity of data produced. Millions of images can be generated from a single survey, a large proportion of which may be ‘empty’ (i.e. not containing any animals), as a result of moving vegetation and changes in light levels which falsely trigger the infrared sensors. This is particularly evident where camera locations are randomly selected and not designed to maximise encounters with the target species [5].

To help analyse such data, a number of software packages have been developed by researchers across the world to automatically classify images from camera traps ([6], [7], [8], [9]). These have proved successful, both at simply removing empty images [10] and at identifying animal species [11] and can achieve similar levels of accuracy to classification by human experts [12].

However, even with the major advances in computer vision technology in recent years, transferability is still a major issue [13]. Many of the recent algorithms [13], such as those presented in [6], [7] and [8], are based on deep-learning methods which, while fast and powerful, mean that their usefulness is contingent on having good-quality training datasets [13]. With camera-trapping becoming more prevalent in a number of habitats [14], obtaining and classifying such training data, which can amount to millions of images [15], may require a large amount of manual classification for each location, completed either by the researchers [16], or with the help of the public [17], something which may be unachievable for projects with lower budgets or lower profiles.

A further barrier for a number of camera-trapping researchers is their coding ability and access to computer resources. There is a wide range of open-source camera-trapping software available, but in general, these require some coding background to install and run [4]. Moreover, deep-learning methods often require expensive hardware, such as GPUs (graphics processing units), to run [10] and hence are unusable for many smaller projects.

The goal of this study was to address these issues through the development of a flexible, low-resource algorithm, Sherlock, that is accessible to researchers with little or no computing background. This algorithm attempts to identify empty images - that is, images that were triggered by an inconsequential movement, such as a plant blowing in the wind. The final product is available on Github [18] and contains simple, detailed instructions on how to use it, including a guide to installing Python.

Sherlock does not require any training data but instead uses an intuitive approach, not contingent on any properties of the habitat in which the images were taken. It examines the sequence of images and looks for marked deviations from a rolling background image, allowing it to remove a large proportion of the false positives from the data. It also does not require large amounts of computing power or a GPU - indeed, it runs off a single core - meaning its hardware requirements are minimal, and that the computer it is running on can continue to be used for other tasks. Moreover, it can be easily paused and restarted, meaning that there is no requirement for it to be continuously run until completion.

Other authors have sought to produce algorithms that do not require any training data, such as [19]. The most similar algorithm that the authors have found to Sherlock is the Zilong algorithm given in [10]. Zilong is founded on similar principles - it does not use neural networks but instead seeks to compare images to the background and is shown to perform well on a range of images in [10]. However, Zilong, alongside the algorithm presented in [19], requires cameras to take “bursts” of images - that is, a single movement triggers a sequence (of at least a certain, fixed length) of images to be taken. However, this assumption was not met in the data on which our algorithm was tested as many encounters between animals and cameras would cause only a single image to be recorded. Moreover, Zilong is more difficult to set up than Sherlock, and so we believe we offer an valuable alternative to a wide range of researchers.

This paper is structured as follows. In Section 2, we give an explanation of the method behind Sherlock, illustrated with an example from our dataset, and also detail the ways in which it can be parameterised. In Section 3, we present the results of testing Sherlock on a set of camera traps positioned across Cornwall, in the south-west of England. Finally, in Section 4, we discuss these results and detail future improvements that could be made to the algorithm.

## 3 Materials and methods

### 3.1 Test data

We tested the performance of Sherlock against a dataset of camera trap images that had been separately classified by one of the authors (VM). Images were obtained from two camera surveys conducted in 2019 in Cornwall, UK, as part of a study investigating analytical methods of estimating badger density. We used a random sample of 23 camera deployments from each site, totalling 240 596 images. Images were classified by a human as either containing a badger or not and were tagged using XnView MP [20]. The dataset contained 306 badger images, with the vast majority being non-badger images but potentially containing other moving subjects such as wildlife, livestock, people and vehicles. The cameras were placed randomly within the survey area rather than being placed strategically to maximise encounters with the target species, in order to satisfy the assumptions of the “random encounter” model for estimating population density [21]. This means that some of the sites had a large number of disturbances caused by sheep and cattle. We also tested performance with varying model parameters using a smaller dataset of 1000 images from a single camera deployment, including 68 classified badger images.

A selection of images which Sherlock correctly tagged as containing badgers is shown in Figure 1, illustrating the range of habitats in which it was used.

**Figure 1:**
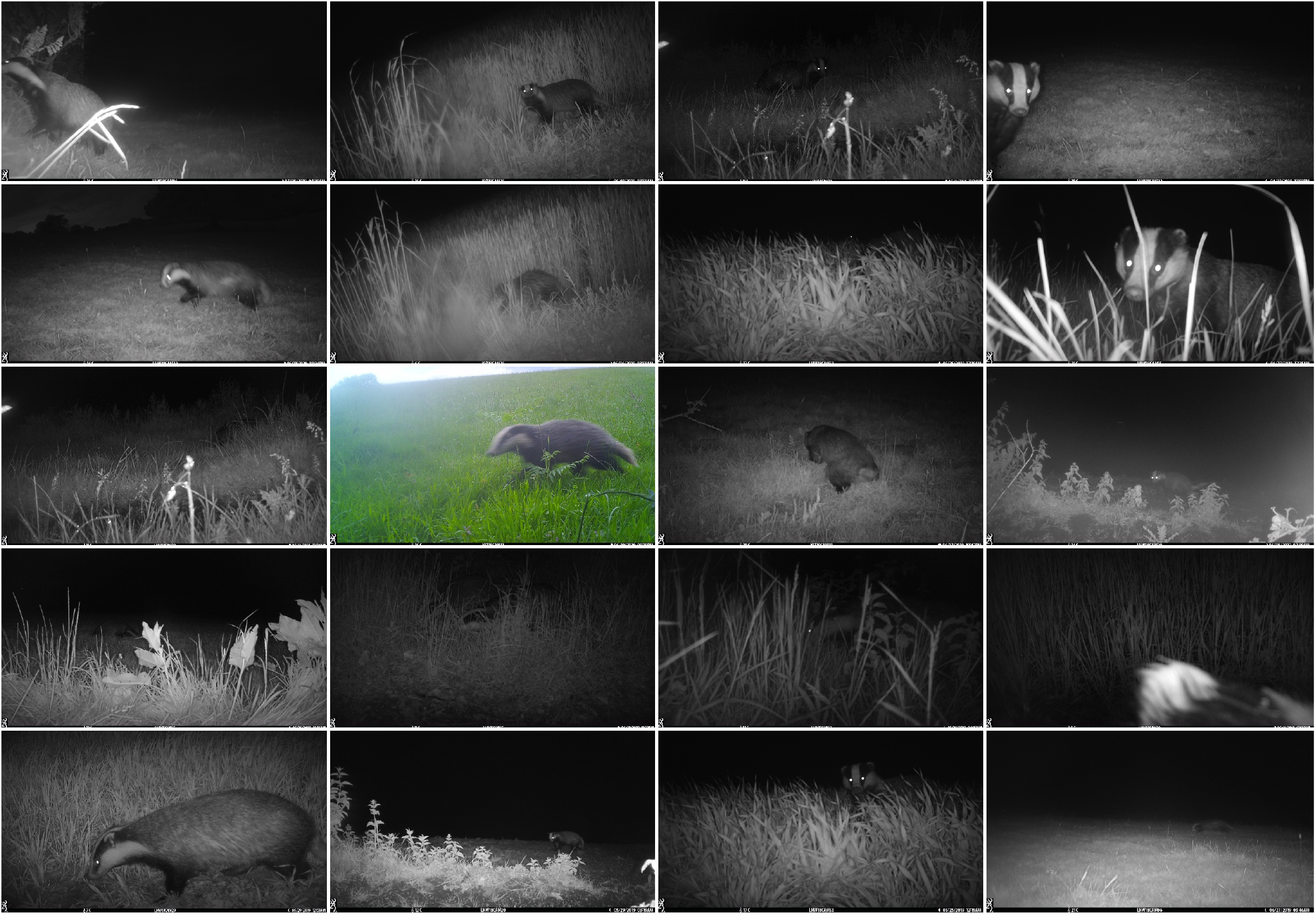
A selection of images which Sherlock correctly identified as containing badgers.

### 3.2 Sherlock code

In this section, the method of the code is detailed, alongside an example. This example is taken from a set of 16 images, shown in Figure 2 which were the only images captured by this camera on the night that they were taken, and hence would have been analysed as a single set by the code (see below for details). The image on which the majority of the analysis is illustrated is image I.

**Figure 2:**
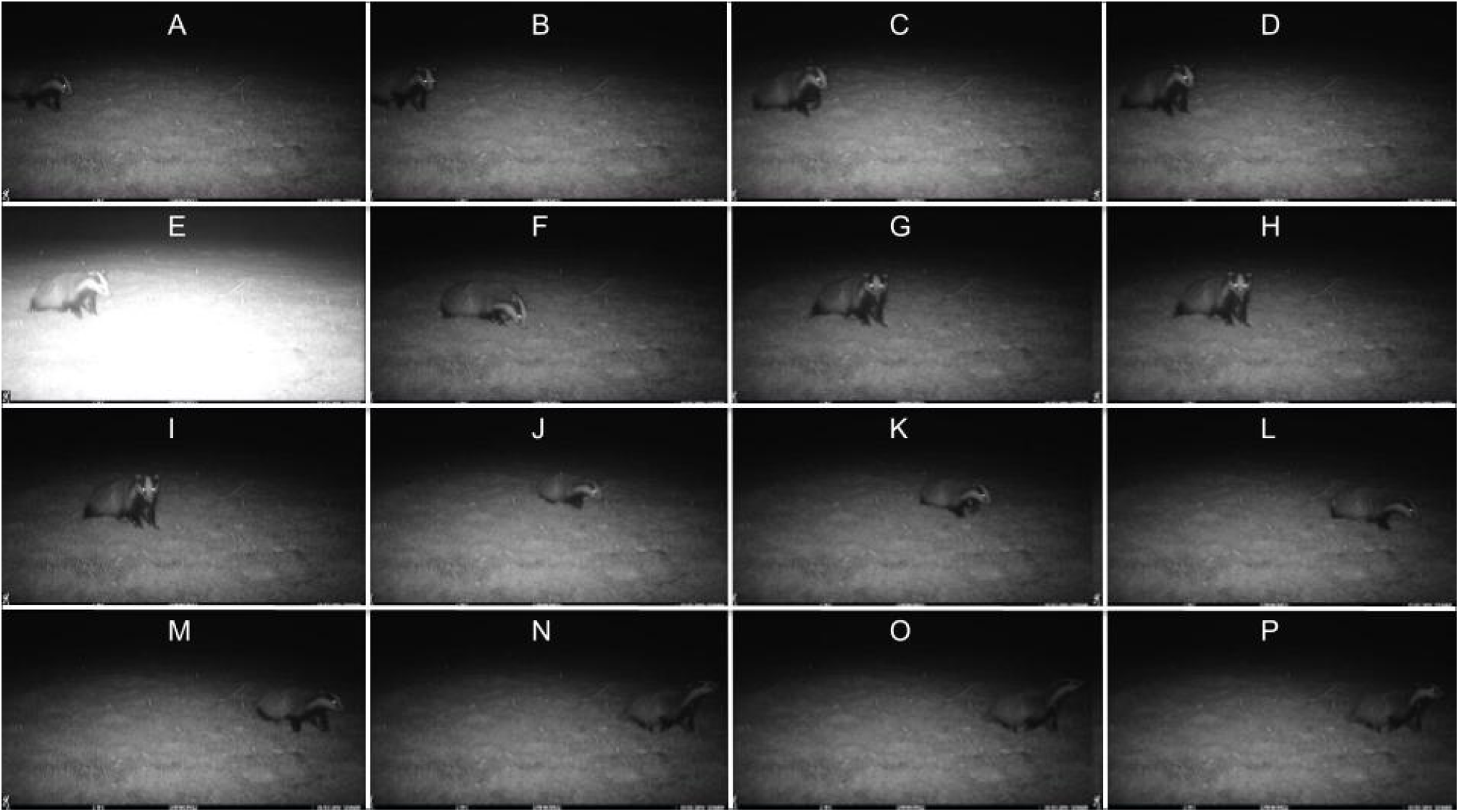
The original set of 16 images which were used to make this example. Although the camera’s infrared flash is automatically adjusted to ensure correct exposure, image E is evidently overexposed. Image I was chosen to illustrate the image analysis procedure

#### 3.2.1 Rolling background image

The main way in which animals are identified is by comparing each image to a rolling background image. Using a rolling background, rather than a fixed one for the whole set of images is particularly helpful for camera traps that are left for a long period of time, where vegetation growth or other factors may change the background. In an ideal setting, this background image would contain the closest *N* images (for some *N*) to the current image. However, this would require computing a separate background image for each image - a computationally expensive task - and instead the background image is periodically updated.

There are two conditions which determine the set of images that comprise each background image. Firstly, there is a user-inputted maximum number of images. This is important as if too many images are used then the background may shift as, for example, light conditions change. Moreover, using a very large number of images may fill up the memory of the computer used and cause the algorithm to crash. The second condition is that the background can only be formed from a set of consecutive daytime images or a set of consecutive nighttime images. Each image is identified as either “day” or “night” by counting the number of pixels that are (approximately) greyscale in the image (images with a high number of greyscale pixels are identified as “night”). This prevents the background image from being a combination of light and dark images and hence ultimately being similar to none of the images from the original set.

Once the set of images has been chosen by the algorithm, the background image is then formed by taking the median colour of each pixel in this set. Using the median instead of the mean prevents large deviations from being over-weighted, giving a more accurate construction of the true background. An example of a background image, created from 16 images, is given in Figure 3, where one can see that the background is recovered, despite the badger walking across the foreground, and the image where the flash caused the foreground to be much lighter. However, the background is not too blurry, as it retains some details such as individual blades of grass, which is an advantage of this background being made from a reasonably small number of images.

**Figure 3:**
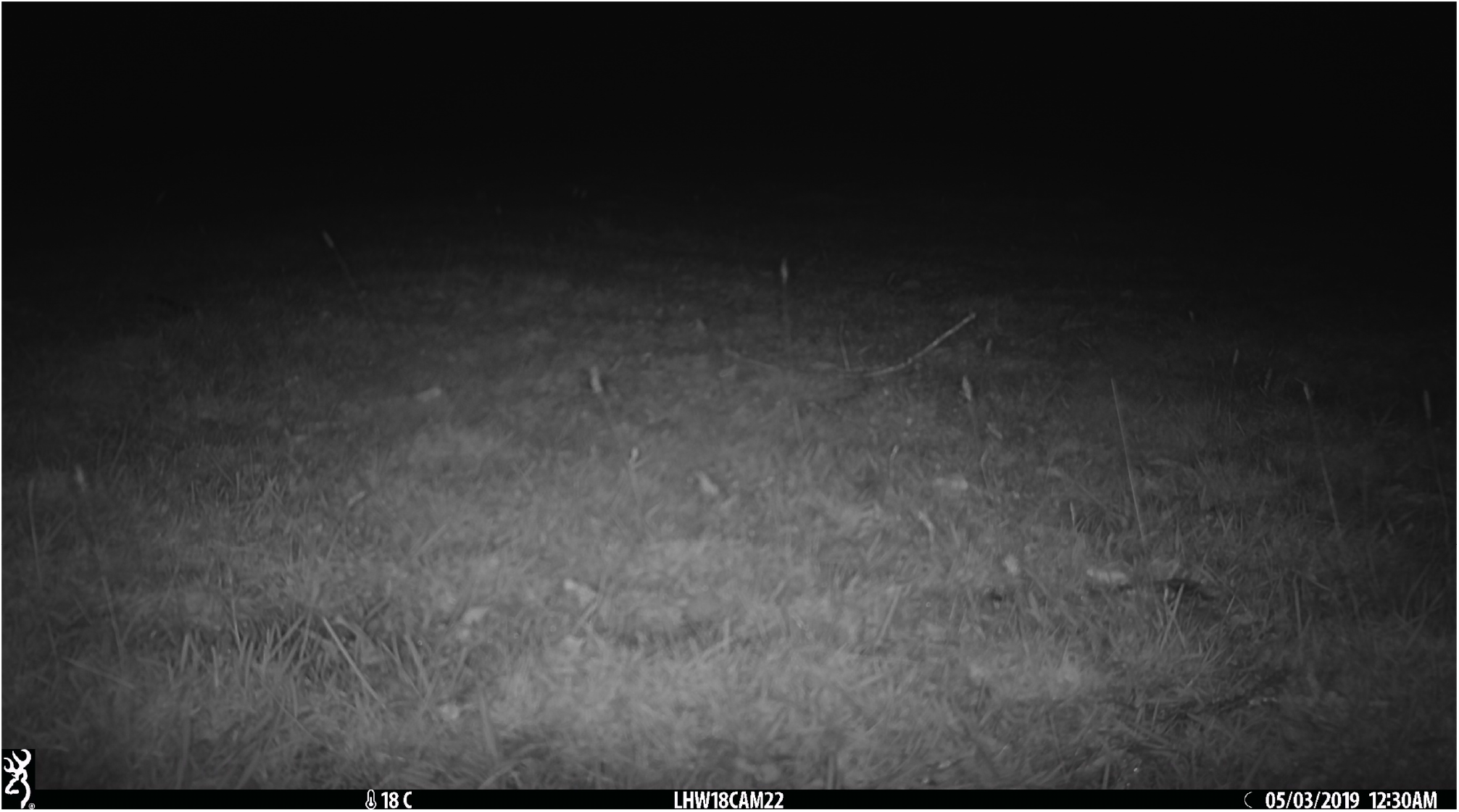
The background image created from the 16 images given in Figure 2

#### 3.2.2 bounding box creation

Once the background image has been calculated, it is possible to test each image and to determine whether it contains an animal. This is done by randomly sampling pixels, testing them and then attempting to build up bounding boxes of “disturbances” - connected sets of pixels that are different from the background image and meet any other criteria (detailed below).

The initial sample of pixels is chosen uniformly (that is, every pixel has an equal chance of being in the sample) at random from the image. The number of pixels sampled can be modified by the user. To determine whether this pixel is a “disturbance”, it must satisfy the following conditions:

1. It must be sufficiently different from the equivalent pixel in the background image (that is, the maximum difference between the RGB values (Red, Green, Blue; with each value being between 0 and 255) of these two pixels must exceed some threshold, which can be different for day and night images)
2. If appropriate, the RGB colours of the pixel must fall between a user-specified upper and lower bound. This condition can be removed by setting the lower bound to (–1, –1, –1) and the upper bound to (256, 256, 256), and was not used by the algorithm when looking for badgers. However, if the animal in question was (at least largely) a single colour, then this parameter could be set so that only pixels close to that colour were considered.
3. If appropriate, the pixel in the image must be sufficiently grey (that is, the R, G and B values of the pixel must have a range smaller than some threshold). This condition can be removed by setting the threshold to be 256, as in many cases (when the species of interest is not approximately greyscale), it will not be appropriate.

Figure 4 shows the results of carrying out this sampling procedure on an image. Most of the disturbances found were indeed on the body of a badger, although there is a minority of small clusters that were caused by moving vegetation or changing light conditions. It is also worth highlighting the number of pixels on the badger that were *not* counted as disturbances, and this noisy feature means that the image requires subsequent analysis

**Figure 4:**
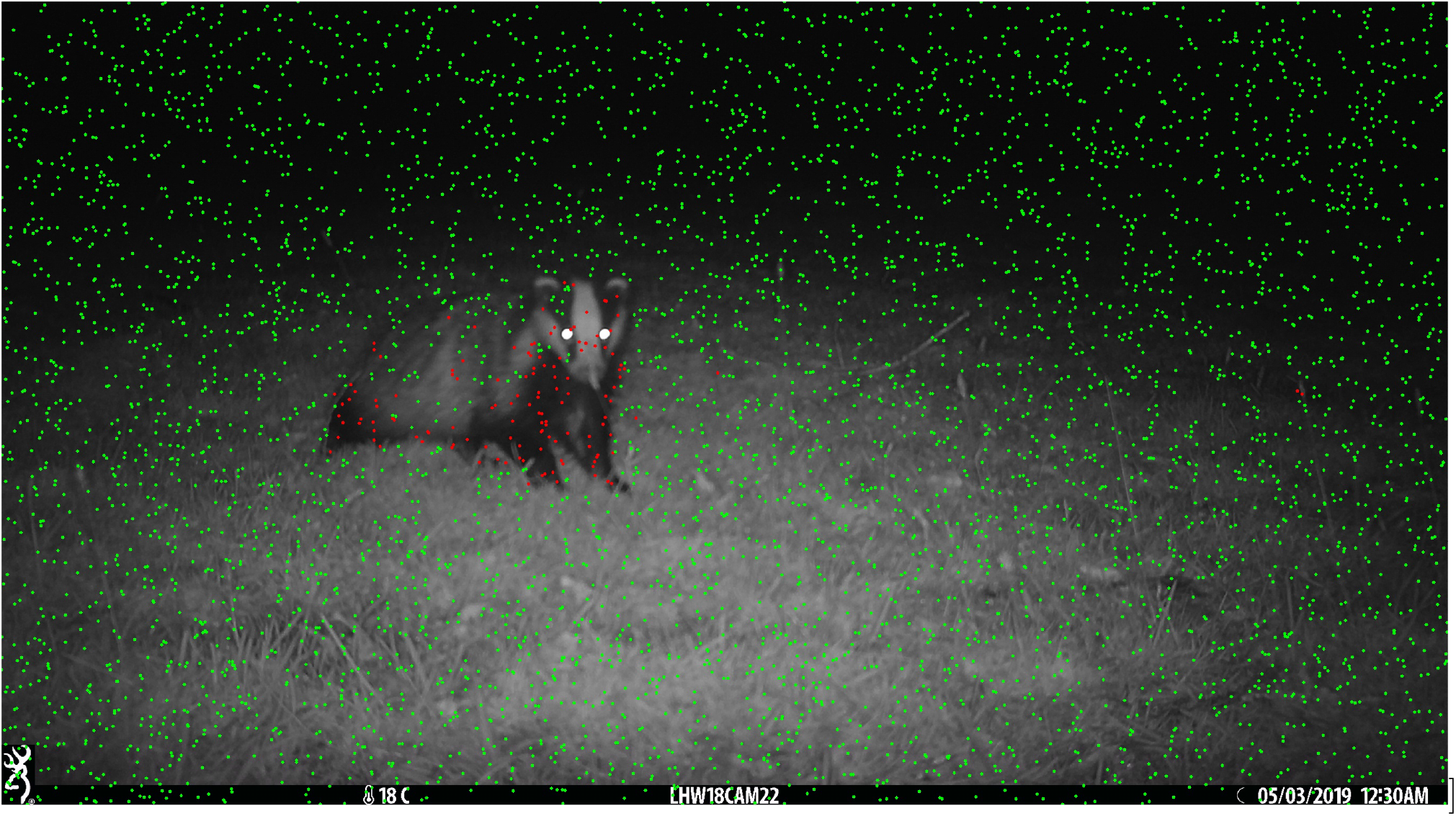
A graphic showing the result of sampling 5000 random pixels image I in Figure 2. Green circles indicate that a pixel was not counted as a disturbance, while red pixels indicate that a pixel was counted as a disturbance. Note that the circles have been set to have a radius of 3 pixels for demonstrative purposes although only a single pixel was analysed in each case.

The area around each pixel that is found to be a disturbance is then analysed further, through what we call the “bouncing” procedure. The algorithm attempts to find paths of pixels moving outwards from the disturbance, as illustrated in Figure 5. It tries four initial directions (upwards, downwards, leftwards and rightwards) in turn, continuing the path in that direction if the next pixel is also a disturbance. When it reaches a point which is not a disturbance, it attempts to change direction (that is, the path “bounces”) to one of the eight horizontal, vertical and diagonal directions, and continues moving. However, it will only try directions which still make progress in the original direction (so, for example, if it was initially moving to the left, it will only try to move diagonally upwards and leftwards or diagonally downwards and leftwards). This process is then continued until there are no more disturbances in any of the allowable directions. The code also allows flexibility for additional bounces to be carried out for each point (with a starting position randomly chosen from within the current bounding box) to improve the accuracy of the bounding boxes created.

**Figure 5:**
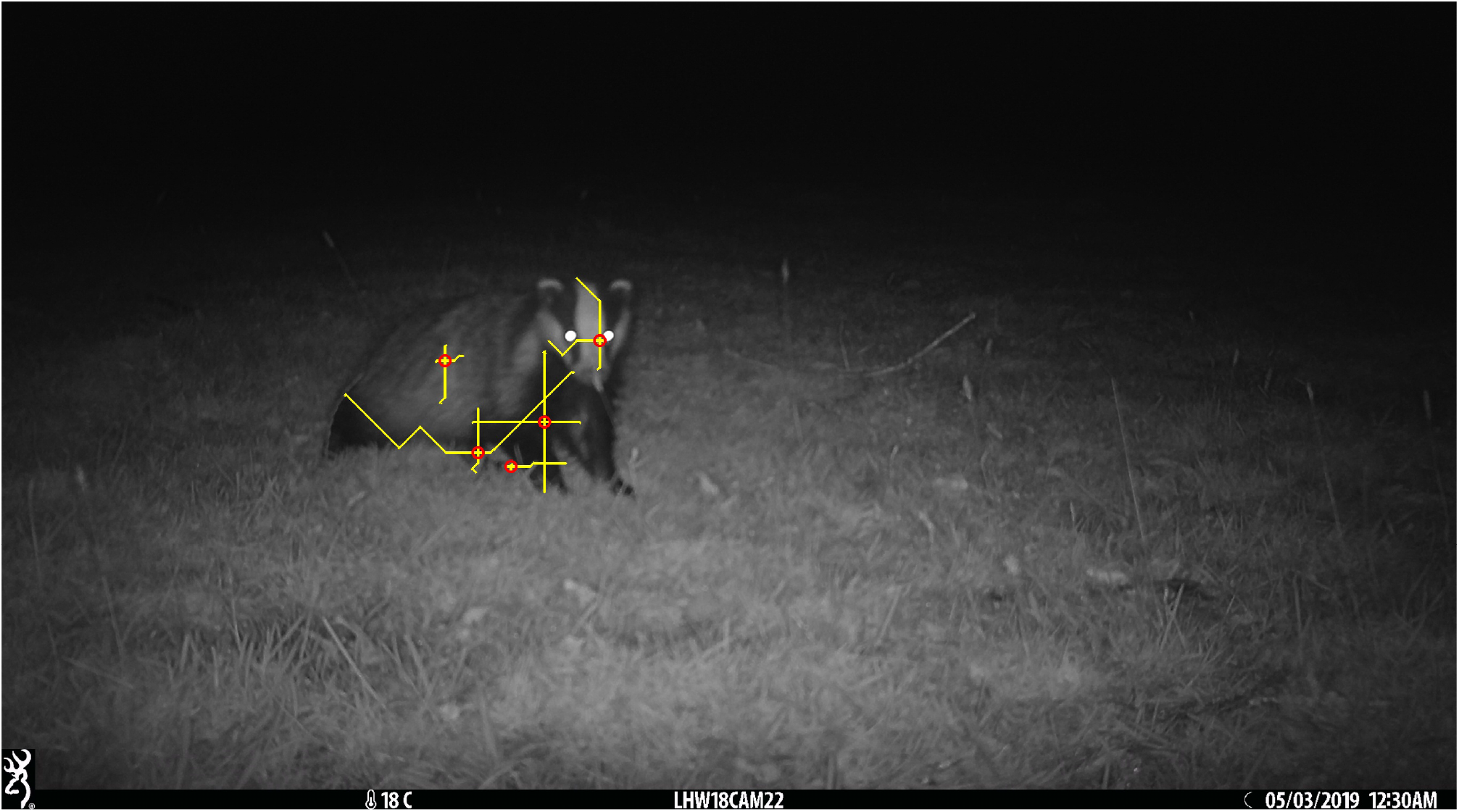
An illustration of five realisations of the “bouncing” procedure. The starting points are shown by red circles, and the bouncing paths are shown by yellow lines. Note that the lines are of width four pixels to improve their visibility, although each considers only a single pixel at a time.

After the bouncing procedure has finished, the maximal and minimal x and y coordinates on the bouncing paths are recorded. Under the assumption that the animal will be an approximately rectangular bounding box, one can then create a bounding box for each path from these maximal and minimal coordinates. If any paths started at separate points have overlapping bounding boxes, these bounding boxes are joined together. The effectiveness of this can be seen in the example of Figure 5 - here, the bounding boxes arising from the paths starting at the five points shown in the figure would be combined into a single bounding boxes.

The main benefit of this novel method is its efficiency. By reducing the problem of finding a two dimensional bounding box to searching for one dimensional paths, the computational cost of the bouncing procedure scales linearly with bounding box size, meaning that the code can run quickly even when a substantial proportion of the image is a disturbance. Moreover, as is shown in Figure 5, one can recover a good approximation to the bounding box from only a small number of bounces, and hence not all of the disturbance points need to be tested - if they are in an already-formed bounding box by the time it is their “turn” to be tested, then the bounce procedure is not carried out.

#### 3.2.3 Bounding box analysis

The final stage of the algorithm analyses each bounding box to determine whether it is likely to be an animal, or simply an inconsequential disturbance.

The first and most important test is the size of the bounding box (that is, its area), which must be larger than a certain threshold (which can be inputted by the user). Different values of this threshold can be used depending on whether the image was a daytime or a nighttime image, to account for the fact that animals are more similar to the background (and therefore likely to create smaller bounding boxes) at night. However, it is important to note that the size of the animals in the image will vary depending on how far away from the camera the animal is, and that the bounding box size should be chosen sufficiently small that all of these animals will be registered.

The second condition is that the bounding box must contain above a certain threshold of pixels that qualify as disturbances. This can help to remove any large rectangular bounding boxes that are created by small objects which have a diagonal orientation (with respect to the image). Moreover, it can help to remove animals whose colouring is, in the majority, outside the colour and greyscale bounds that have been specified by the user (if, for example, the user is only interested in specific species).

#### 3.2.4 Adjacent images

It is often the case that images containing animals occur in sequences (as the animal moves through the field of view of the camera). To account for this, one can choose a value *n* such that if an image is within n images of an image that has been classed as a positive, and the timestamp of this image differs by at most some threshold (set as default to 20 seconds) from the timestamp of the positive image, then it is also classed as a positive, irrespective of what its classification would have been otherwise. We call these images adjacent images.

Note that adjacent images that would otherwise have been classed as negatives do not trigger their own set of additional positive images. This step is helpful as animals often trigger a long sequence of images, some of which may be misclassified by the algorithm, and this step helps to find these errors. The default value of *n* is 1, which strikes a good balance between increasing sensitivity without decreasing specificity to a large degree.

#### 3.2.5 Extending the set of positive images

An additional consideration is that images containing animals can affect the classification of those images near to them because they may interfere with the background image created, particularly if an animal moves close to the camera. This will be a problem regardless of the difference in times between this image and the images near it in the image sequence. Thus, we consider “extending” the set of positive images by including the images either side of each image originally in the set of positives (including adjacent images). In the results section, we present separately the cases where the dataset is and is not extended.

#### 3.2.6 Algorithm outputs

Sherlock puts images in three categories - those which potentially contain the animal of interest, those which it believes do not contain the animal of interest, and images that caused an error when being processed (this is very rare, but may occur if an image has been corrupted). The basic summary of this categorisation is found in CSV (comma separated value) files which it outputs into each folder that contains images. Secondarily, Sherlock can create a new copy of each image, with the addition of a red box drawn around anything it believes to be an animal of interest, in a folder based on its categorisation. This may be of use if the potential animal images are then categorised by a human, as it will highlight which disturbances Sherlock has categorised as an animal.

### 3.3 Code runtime

The test dataset was processed by Sherlock on a Dell laptop with a clock speed of 2.11GHz and a RAM of 32.0GB. The images were stored on an external hard drive, and each image was a JPEG file between 1 and 2 megabytes in size, containing just over 4 million pixels. The code was run through the Python 3 user interface, Jupyter Lab. The 240 596 images in the test dataset were processed in approximately a week, which equates to roughly 5 seconds per image. The code was run on a single core, but could be easily parallelised - in particular, images from different sites are processed entirely independently.

Two smaller tests were carried out by the authors KLA and CH where only 1000 images were processed. These were performed on laptops with similar specifications to the above.

## 4 Results

### 4.1 Overall results

Table 1 shows that Sherlock achieved a reasonably good performance, removing close to 50% of the images in the unextended set. Moreover, the fact that only 8.1% of the images and 4.5% of the sequences were missed in the two cases shows that Sherlock’s ability to reduce the need for human tagging comes with only a small cost in the reduction of relevant data.

**Table 1:** Sherlock’s performance on the test data. Both the cases when the set of positive images was extended (that is, images which were immediately before or after an image classified as a positive were counted as positives) and when the set of positive images was not extended are considered.

| Classification method | Number (% correctly classified) of: |  |  |
| --- | --- | --- | --- |
|  | Images |  | Sequences |
|  | With badgers | Without badgers | With badgers |
| Human | 308 | 240288 | 67 |
| Sherlock (unextended) | 283 (91.9%) | 118351 (49.3%) | 64 (95.5%) |
| Sherlock (extended) | 302 (98.1%) | 100235 (41.7%) | 66 (98.5%) |

Depending on the exact parameters of the study, it may be beneficial to extend the set of positives. In this case, as one would expect, the percentage of false positives increased from 50.7% to 58.3%. However, almost all (98.1%) of the badger images were correctly identified in this case, highlighting the fact that many of the false negatives from Sherlock were simply a part of a sequence of images of badgers. The fact that only one sequence was missed in this case highlights Sherlock’s sensitivity, and suggests that researchers can be confident that the vast majority of the useful data will be retained.

### 4.2 Performance by site

Part of the reason for Sherlock’s low precision is that many of the sites on which it was tested contained a number of other animals which were not badgers, but which had similar coloration (such as cattle and sheep). Table 2 shows that Sherlock performed poorly on sites where sheep and cattle are abundant, but performed markedly better on sites where no livestock were present. Indeed, the false positive rate of 7.7% is very similar to the 7.5% of Zilong [10] and shows that Sherlock could be an extremely useful tool on similarly clean datasets.

**Table 2:**
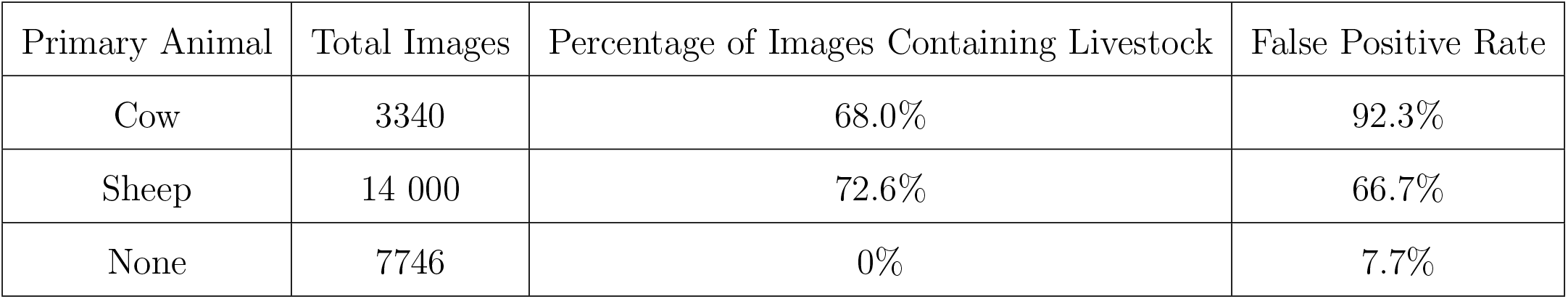
Sherlock’s performance across three pairs of sites, with the majority of images from the first pair being of cattle, the majority in the second pair being of sheep, and all of the third pair containing no livestock. Across all sites, 23 of the images contained badgers, of which 20 were correctly identified. Note that these results are based on the unextended set of positives.

| Primary Animal | Total Images | Percentage of Images Containing Livestock | False Positive Rate |
| --- | --- | --- | --- |
| Cow | 3340 | 68.0% | 92.3% |
| Sheep | 14 000 | 72.6% | 66.7% |
| None | 7746 | 0% | 7.7% |

**Table 3:** The results of using different parameter values on a test dataset containing 1000 images, 68 of which contained badgers. Note that these results are based on the unextended set of positives.

| Bounces | Pixel Sample Size | Run time (minutes) | Precision | Recall |
| --- | --- | --- | --- | --- |
| 4 | 5000 | 31.1 | 15% | 93% |
| 4 | 1000 | 16.0 | 16% | 91% |
| 20 | 5000 | 67.0 | 15% | 93% |
| 20 | 1000 | 34.0 | 14% | 84% |

### 4.3 Testing with other parameter sets

There was, in general, very little difference between the precision and recall achieved by these algorithms, though the algorithms with smaller sample sizes did have a slightly recall. Most surprising was the drop in precision of the algorithm with a sample size of 1000 and 20 bounces, which performed worse than the algorithm with a sample size of 1000 and 4 bounces. This could have been caused by the increase in randomness in the results which occurs when a smaller sample size is used. It also suggests that there may be little value in increasing the bounce parameter, particularly as it comes with the cost of increased runtime. Indeed, setting this parameter too high could lead to bounding boxes which should be separate being joined together.

The clearest difference between the four parameter sets was in the runtime of the algorithm, with the quickest performing around 75% quicker than the slowest, and with only a small decrease in sensitivity. Further investigation, and a proper definition of optimality would be needed to determine the “optimal” set of parameters, but it certainly suggests that there may be scope to speed up the algorithm, particularly if one is willing to accept a small drop in sensitivity.

### 4.4 Comparison to human tagging

**Table 4:** The results from two of the authors separately tagging the same set set of 6531 images with and without the use of Sherlock. Note that, in addition to the human tagging time, Sherlock took between 3 and 4 hours to process these images on each of the authors’ computers.

| Tagger | With/without Sherlock | Tagging time (minutes) | Badgers tagged | Sequences |
| --- | --- | --- | --- | --- |
| CH | Without | 20.9 | 23 | 6 |
| CH | With | 13.3 | 22 | 6 |
| KLA | Without | 25.8 | 20 | 6 |
| KLA | With | 7.8 | 21 | 6 |

To assess the practical usefulness of Sherlock, two of the authors (KLA and CH) separately tagged the set of 6531 images with and without the use of Sherlock. The results are summarised in Table 4.

In both cases, Sherlock substantially reduced the human time taken to tag these images. Moreover, Sherlock had only a small effect on the number of images that each person tagged as containing a badger and, in one case, this effect was positive as KLA found an additional image that contained a badger. Indeed, the largest source of error in this process appears to be the human doing the tagging as, given an identical set of images without Sherlock, CH tagged 15% more images as badgers than KLA.

In both cases, Sherlock took between 3 and 4 hours to process these images. While this is substantially longer than the human tagging time, researchers were of course free to do other work while Sherlock was running and thus it can still greatly increase the efficiency of processing camera trap data.

## 5 Discussion

Sherlock has the potential to substantially reduce the number of human hours required to process a camera-trapping dataset, as it can remove a large proportion of unwanted images while losing very few images of interest. It has performed well across a range of camera placements which contained many complicating factors such as vegetation and other animals that were not of interest. Moreover, an almost identical algorithm has been used by one of the authors (MJP) in their work for Oxford City Football Club for tracking players from video footage of their matches, highlighting the algorithm’s flexibility and the potential for its widespread adoption. It is simple to set up, with detailed instructions available on our GitHub page [18], and does not require high levels of computing power to process images at a reasonably quick rate.

Across the whole dataset, Sherlock did not achieve as high a specificity as some similar algorithms. For example, Zilong [10] removed approximately 92.5% of the empty images from its dataset, compared to the 49.3% that Sherlock achieved from the unextended set of positives. However, such a direct comparison may not be an appropriate assessment of the relative utility of the two algorithms. When sites with no livestock were considered, Sherlock was able to remove 92.3% of the images, which is very similar to Zilong’s performance. This suggests that Sherlock’s apparent low performance was simply due to the high levels of noise in the data, and that it has the potential to be an extremely useful image-removal tool, particularly when the camera trap has little interference from other animals, or when all animals are of interest.

Moreover, the fact that Sherlock has such a high sensitivity of 91.9% - higher than, for example, the reported 87% of Zilong - means that it could be used as a tool to pre-process a large set of data, reducing the number of images that need to be classified by more computationally intensive algorithms that, for example, might seek to classify animals to species level. The high sensitivity means that very little useful information would be lost during this step, while the computational savings may be valuable, both in terms of financial and time cost. This would be particularly relevant in the examples considered in Table 2, where, in the sites with sheep, Sherlock was able to remove a reasonable proportion of images from a very noisy dataset. There are a number of ways in which Sherlock could be refined to increase its performance. Primarily, adding the option to parallelise the algorithm, and in particular adding the option to run it on a GPU, would lead to a large reduction in the amount of time taken to process each image (at the expense of making it less accessible). Moreover, it may be possible to more intelligently sample pixels from the image, so that the perspective of the image is taken into account - one needs to sample fewer pixels in the foreground than the background to achieve the same level of coverage. Combining this with position-dependent-bounding box-size restrictions could further improve performance, particularly if there is vegetation close to the camera that frequently triggers it.

There are also some limitations to the method of forming the background images. Firstly, there is not always an immediate transition between daytime and nighttime images, and so there may be some images taken at dawn or dusk which are very dissimilar to the background. These images will then be classified as potentially having an animal, adding to the number of false positives in the dataset. However, this was true for a small proportion of the images and so did not substantially affect algorithm performance.

The second limitation is that an animal that comes very close to the camera may cause a daytime image to be misclassified as a nighttime image (as the resultant image may be almost entirely black). To combat this, if there are fewer than 3 images in a single day or night period, the code will assign them as having an animal. Again, while this adds to the number of false positives, it happens sufficiently rarely that performance is not substantially affected.

A final limitation is that a habitat where the background is naturally grey may lead to a number of images being misclassified as being nighttime images. However, such environments (such as rocky mountains, or cities) are not commonly used for camera trapping.

Despite these limitations, we believe that, in its current form, Sherlock has the potential to be of use to a wide range of researchers. The levels of specificity and sensitivity that it achieves can substantially reduce the amount of processing by humans required in camera-trapping projects Moreover, its ease of use and minimal hardware requirements mean that it is available to the vast majority of researchers, and we hope that its adoption can help accelerate many areas of biological research.

## 7 Author Contributions

Conceptualization: MJP, VM Data Curation: VM Investigation: VM, KLA, CH Methodology: MJP Software: MJP Validation: VM Writing - Original Draft: MJP, VM Writing - Review and Editing: MJP, VM, RW, MR, CAD

## 8 Data Availability

The code for Sherlock is currently available on GitHub, and will be archived on Zenodo upon publication of this article. The underlying image dataset will be also be archived on Zenodo.

